# Plant Bioindicators of Pollution in Sadat City, Western Nile Delta, Egypt

**DOI:** 10.1101/856229

**Authors:** M. F. Azzazy

## Abstract

This study investigated using plants growing in industrial and residential areas as bioindicators and biomarkers of industrial pollution by analyzing flavonoids and metals in *Bougainvillea glabra* leaves by HPLC-MS, neutron activation analysis, and atomic absorption spectrophotometry. There were significantly higher levels of flavonoids and phenolic compounds in plants growing in industrial areas compared to those growing in residential zones (P<0.05). Metal accumulation in leaves was also significantly higher in the industrial zone than the residential zone: iron, lead, zinc, nickel, and manganese were present at significantly higher levels in plants growing near steel factories compared to those growing in the residential zone (P<0.05). Air, water, and soil samples associated with the studied plants in both areas were analyzed for metals. In general, heavy metal concentrations in the industrial zone were significantly higher than those in the residential zone, but the concentrations of the heavy metals in air, water, and soil were under legal environmental limits.

## 1. INTRODUCTION

Air- and water-borne pollution is an undesirable consequence of industrialization that is having a growing impact on productivity, health, and climate change [1–5]. Pollution affects plants and animals via a number of routes including through pollutants dissolved in the rain (e.g., sulfur dioxide producing sulfuric acid), chemical discharge into water courses, and particulate matter (PM) in the air including dust, dirt, and smoke. Pollution impairs plant photosystem II [1] and is associated with respiratory and cardiovascular diseases [3]. Monitoring and assessing pollution in the air, water, and soil from industrial sources is an important aspect of managing the impact of industry and pollution on our environment [4]. Bioindicators are living organisms such as plants, planktons, animals, and microbes with the capacity to monitor the health of the environment. They are therefore a useful means to measure the negative impacts of industrial activity on the environment [6]. Similarly, environmental biomarkers are identifiable measures (e.g., chemicals or genes) of environmental processes [7] and a useful tool not only to monitor and evaluate the environmental state but also develop our knowledge of molecular toxicity mechanisms in different animal and plant species in the ecosystem [8].

Therefore, plants can be used to assess whether certain ecophysiological responses may be useful biomarkers of urban pollution. Some plant biomarkers are specific to only one pollutant or group of pollutants, while others respond to a wide range of pollutants and/or stressors [9]. For example, flavonoids and phenolic compounds used to indicate stress in plants [10,11]. These secondary metabolites, in particular polyphenols, are of particular importance in plant-environment relationships [12]

*Bougainvillea glabra* (family: Nyctaginaceae) is a common ornamental plant grown in tropical and subtropical gardens and grown as a shrub or climber [13]. The aim of the present study was to investigate using *B. glabra* as a bioindicator and assess flavonoid, phenolic compound, and metal accumulation in their leaves as biomarkers of environmental pollution.

## 2. MATERIALS AND METHODS

### 2.1 Study area

Sadat City is located about 100 km north of Cairo, Egypt. The city is bounded 30°19’30”-30°40’27” E longitude and 30°15’50”-30°34’00”N latitude (**Fig. 1**), from the east by Kafer Dawoud and El Khatataba, from the west by El Birigat, and from the north by Nubariya Canal and El Tahrir. Industrial zones are located in a separate spine along the south-eastern edge of the city to ensure that industrial pollution travels downwind. To protect the city from wind and storms, a green shelterbelt of trees about 35,000 feddans (1 feddan = about 175 m^2^) in area was planted around the city, containing about 2,000 feddans of vegetables and fruit.

**Fig.1.**
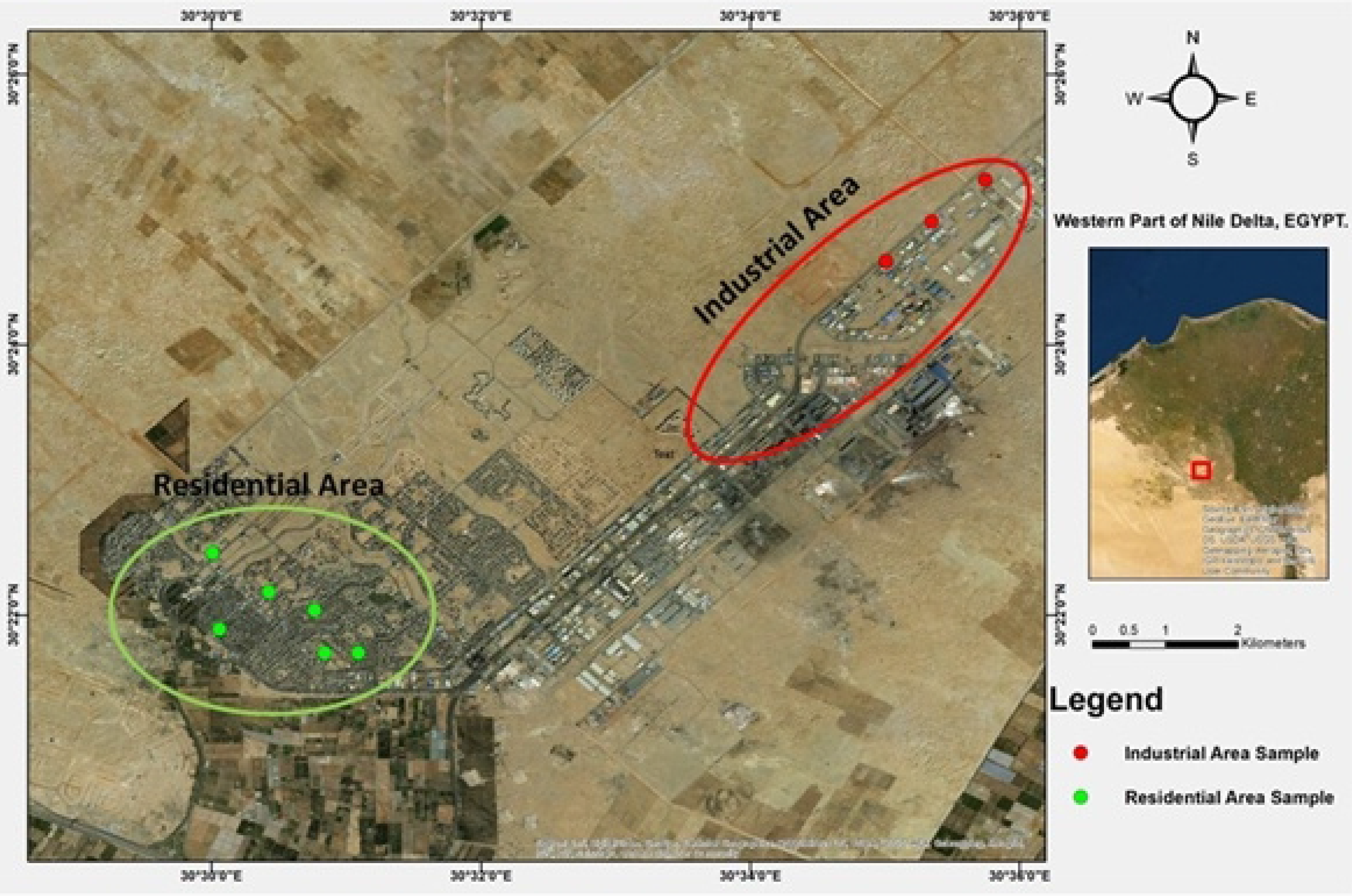
Map showing the sites of sample collection in the study area.

The residential districts and industrial zones of Sadat cover about 50 km^2^ of the study area. With a future planned capacity of 750,000 inhabitants, the city currently hosts a population of about 200,000 inhabitants and about 100,000 seasonal residents working in industry and agriculture. The city has one waste disposal site at the southern side of the city behind the industrial zone about 700 feddans in area.

### 2.2 Sampling

The study area was sampled between January 2018 and December 2018 at the sites shown in **Fig. 1**: six in the residential area and three at three industrial sites (El Dawlia Oil, El-Ezz Steel, and Prema Ceramics). PM_10_ (particulate matter <10 micrometers) and heavy metals were sampled along with the physical and chemical properties and heavy metal content of soil, water, industrial chimneys, and inside the factories. Plants were studied morphologically. Flavonoids and phenolic compounds were investigated in *B. glabra* leaves by HPLC-MS, while metals were measured in leaves using neutron activation analysis and atomic absorption spectrophotometry.

### 2.3 Climatic parameters

The Tropical Rain Measurement Mission (TRMM) records three hourly rainfall measurements and is freely available online including from the TOVAS website (https://serc.carleton.edu/resources/22961.html). Climate was also estimated using Egyptian Meteorological Authority data between January and December 2018.

### 2.4 Identification of the collected plants in the study area

Plants in industrial and residential areas were identified using well identified herbarium specimens of the Environmental Studies and Research Institute (ESRI) and the Herbarium Faculty of Science Botany Department, Mansoura University. A voucher specimen was deposited at the herbarium of the ESRI, University of Sadat City. The morphological features of plants growing in both locations were compared.

### 2.5 Biomarker analysis

HPLC-MS Ultimate 3000/Amazon SL ion trap mass spectrometry was undertaken using Acclaim 2.2 μm 120 A 2.1-150 mm columns (Thermo Fisher Scientific, Waltham, MA). The HPLC-MS device was used to detect flavonoids and phenolic compounds in plant leaves.

### 2.6 Chemicals and reagents

Methanol, acetonitrile, and deionized water were of HPLC-MS grade and were from Sigma Aldrich (St Louis, MO).

### 2.7 Plant materials and sample preparation

Leaves were collected from plants growing in the study areas. Methanol extraction was performed with a Soxhlet apparatus [14]. Extraction efficiency was enhanced by the application of the methods of [15, 16]. Leaves were dried and ground into powder, and 0.1 g of powdered leaves was mixed with 10 ml 70% methanol and placed on a rotating shaker at 200 rpm at 40°C for 16 h. The filtrate was collected and filtered through a 0.45 μm nylon filter.

A volume of plant extract was injected into the HPLC-MS column for analysis. All sample solutions were stored at 4°C prior to use. The compound at each peak was recognized according to its molecular weight and retention time.

### 2.8 Determination of phenolic contents

The total phenolic content was determined for plant leaves extract using the Folin–Ciocalteu method [17]. Briefly, 1 mL of extract (100–500 g/mL) solution was mixed with 2.5 mL of 10% (w/v) Folin–Ciocalteu reagent. After 5 min, 2.0 mL of Na_2_CO_3_ (75%) was subsequently added to the mixture and incubated at 50°C for 10 min with intermittent agitation. Afterwards, the sample was cooled and the absorbance was measured utilizing a UV spectrophotometer (Shimazu, UV-1800) at 765 nm against a blank without extract. The outcome data were expressed as mg/g of gallic acid equivalents in milligrams per gram (mg GAE/g) of dry extract.

### 2.9 Determination of flavonoid contents

The flavonoid contents of plant leaf extracts were measured as per the Dowd method [18]. An aliquot of 1 mL of extract solution (25–200 g/mL) or quercetin (25–200 g/mL) were mixed with 0.2 mL of 10% (w/v) AlCl_3_ solution in methanol, 0.2 mL (1 M) potassium acetate, and 5.6 mL distilled water. The mixture was incubated for 30 min at room temperature followed by the measurement of absorbance at 415 nm against the blank. The outcome data were expressed as mg/g of quercetin equivalents in milligrams per gram (mg QE/g) of dry extract.

### 2.10 Instrumentation

Plant specimens were studied for flavonoids and phenolics in the central laboratory of the ESRI, University of Sadat City, Egypt. The binary mobile phase consisted of solvents A (1% formic acid) and B (acetonitrile with 1% formic acid). A mobile phase gradient was used as follows: 0 min, A: B 10–90; 36 min, A: B 70–30; 50 min, A: B 100–0. After each run, the chromatographic system was equilibrated with 20% B for 30 min. The injection volume was 20 μl. Flow rate was 0.8 ml/min. The effluent was split in a ratio of 2:3 using a micro-splitter valve before introduction into the mass spectrometer. UV traces were measured at 290, 254, and 350 nm and UV spectra (DAD) were recorded between 190 and 900 nm according to [19].

### 2.11 Gas analysis

NO, NO_2_, and NOx were measured with the Thermo Environmental Instruments 42C NO, NO_2_, NOx Analyzer, EPA reference method RFNA-1289-074, over 10-300 seconds. SO_2_ was measured with the Thermo Environmental Instruments 43C SO_2_ Analyzer, EPA equivalent method EQSA-0486-060, over 10-300 seconds. CO was measured with the Thermo Environmental Instruments 48C CO Analyzer, EPA reference method RFCA-0981-054, over 10-300 seconds. Outdoor NO_2_ concentrations were measured using the method in [20]. Gases were measured in the work environments using Testo Portable Emission Gas Analyzer (Testo Inc., Sparta, NJ; Model 350M/XL).

### 2.12 Particulate matter (PM)

Particulate matter (PM) was collected using the filtration method [21]. Particles less than 10 μm in diameter (PM_10_) were trapped by the filter, and a TSP Meter (Kanomax, Andover, NJ) was used to measure TSP in the work environment and PM_10_ apparatus to measure PM_10_ according [22].

### 2.13 Atomic absorption spectrophotometry

Atomic absorption spectrophotometry was performed using a Spectrum 100 (Perkin Elmer, Waltham, MA) to determine heavy metal concentrations in plants.

### 2.14 Neutron activation analysis

The concentration of sodium (Na), calcium (Ca), magnesium (Mg), chloride (Cl), and manganese (Mn) in plant tissues was assessed by neutron activation analysis (NAA) using the IBR-2 Reactor (IBR2 of the Joint Institute for Nuclear Research, Dubna-Russia (JINR).

### 2.15 Groundwater samples

Two water samples were collected from two wells in the study areas (industrial and residential, one sample from each). The water samples were acidified and stored in an icebox before being analyzed by atomic absorption spectroscopy (AAS) at the Water Department, Central Health Laboratories, Ministry of Health and Population, Egypt.

### 2.16 Soil chemical analysis

Soil samples were collected from the industrial and residential zones (three samples from each), ground, and digested with HF and HClO_4_. One gram of each sample was placed in a Teflon beaker and treated with 10 ml HF and 2 ml HClO_4_ before being heated to nearly dry the sample. The residual material was dissolved with HCl (12 N) and the filter diluted with 25 ml deionized water [23, 24]. Chemical analysis was performed by atomic absorption spectroscopy (AAS) for chromium (Cr), nickel (Ni), and lead (Pb) [23].

### 2.17 Statistical analysis

Data are expressed as mean ± standard deviation (SD). Data were analyzed with t-tests using SPSS ver. 14.0 (IBM Statistics, Chicago, IL). A p-value < 0.05 was considered statistically significant.

## 3. RESULTS

### 3.1 Climatic study

Monthly mean temperatures over the study period varied from a minimum of 12.4°C in January to a maximum of 36.4°C in July over the study period (Table 1). Rainfall ranged from 21.4 mm in January to no rainfall in June, July, and August. Relative humidity ranged from 45.0% in December to 71.0% in August. Wind speed varied from 6.475 m/s in October to 10.36 m/s in March.

**Table 1.** Means of meteorological parameters from January to December 2018

| Month | Air temperature (°C) | Highest of rainfall mm/month | Relative humidity (RH) % | Wind speed (km/h) |
| --- | --- | --- | --- | --- |
| January | 12.4 | 21.4 | 56.0% | 9.435 |
| February | 13.5 | 15.2 | 57.0%% | 9.99 |
| March | 15.7 | 16.7 | 60.0% | 10.36 |
| April | 19.3 | 16.8 | 62.0% | 9.99 |
| May | 22.7 | 15.9 | 66.0% | 7.955 |
| June | 25.8 | Trace | 68.0% | 8.325 |
| July | 36.4 | Trace | 70.0% | 7.215 |
| August | 37.8 | Trace | 71.0% | 6.66 |
| September | 25.0 | 2.6 | 70.0% | 7.955 |
| October | 22.1 | 7.1 | 67.0% | 6.475 |
| November | 17.8 | 18.3 | 55.0% | 7.03 |
| December | 13.8 | 16.0 | 45.0% | 7.77 |

#### Morphological changes in plant leaves due to pollution

Overall, the leaves of plants grown in industrial zone had discolored, dusty, and wrinkled leaves compared to their counterparts growing in the residential zone, which had smooth surfaces and edges and were clean with no wrinkles and normal edges (Fig. 2).

**Fig. 2.**
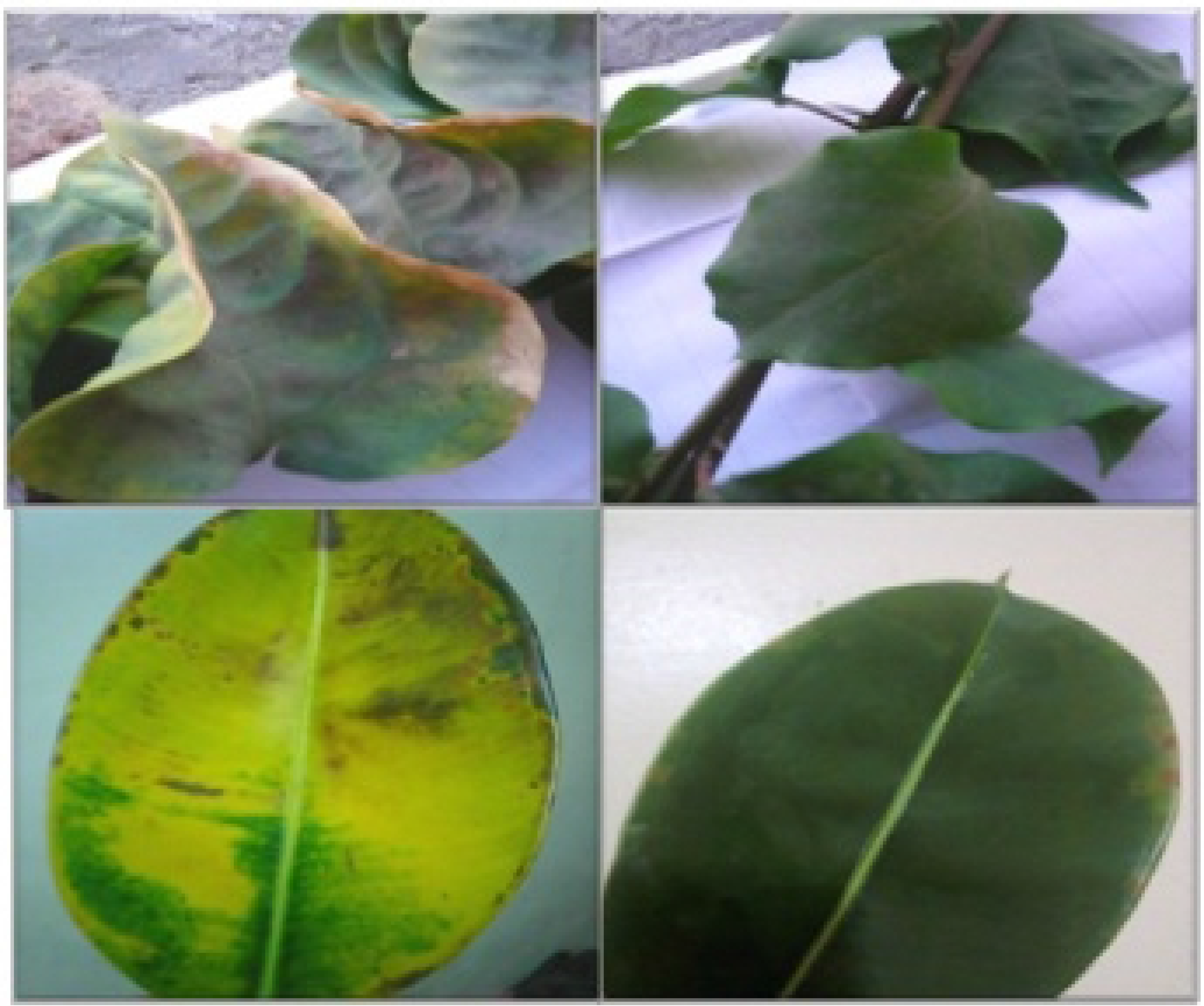
Illustrative morphological differences in *Bougainvillea glabra* growing in industrial (left) and residential (right) zones.

### 3.2 Metal concentrations in plants growing in industrial and residential zones

Table 2 shows metal concentrations in *Bougainvillea glabra* leaves growing in the industrial and residential zones. Iron, zinc, lead, nickel, and manganese were all present at significantly higher levels in plants growing in the industrial zone than the residential zone (P<0.05), while copper and cadmium were undetectable in both.

**Table 2.** Heavy metal content in *Bougainvillea glabra* leaves in industrial and residential zones. NS=not significant, ppm=parts per million; values are means ± SD from three replicates.

| Item | Industrial Zone | Residential Zone | Sign. |
| --- | --- | --- | --- |
| Fe (ppm) | 494.00 ± 1.63 | 0.0 ± 1.63 | P<0.05 |
| Zn (ppm) | 445.00 ± 2.41 | 33.28 ± 2.41 | P<0.05 |
| Pb (ppm) | 0.0066 ± 0.001 | 0.0 ± 0.001 | P<0.05 |
| Ni (ppm) | 74.11 ± 0.10 | 53.31 ± 1.7 | P<0.05 |
| Mn (ppm) | 452.47 ± 3.9 | 211.00 ± 6.3 | P<0.05 |
| Cu (ppm) | 0.00 | 0.00 | NS |
| Cd (ppm) | 0.00 | 0.00 | NS |

### 3.3 Plant flavonoid analysis

HPLC-MS analysis of *B. glabra* (Table 3 and Fig. 3) leaf extracts revealed a set of peaks in plant extracts derived from the industrial zone. Twenty-one compounds were detected in industrial zone leaves; five major peaks were selected on the basis of peaks and retention times (Rt). Rosmarinic acid had an Rt of 48.9 min and peak area percentage of 12.01%; chlorogenic acid had an Rt of 6.7 min and peak area of 3.9%, quinic acid had an Rt of 32.0 min and peak area of 3.8%, R-adrenaline had an Rt of 11.0 min and peak area of 2.9%, while caffeine had an Rt of16.8 min and peak area of 2.5%.

**Table 3.**
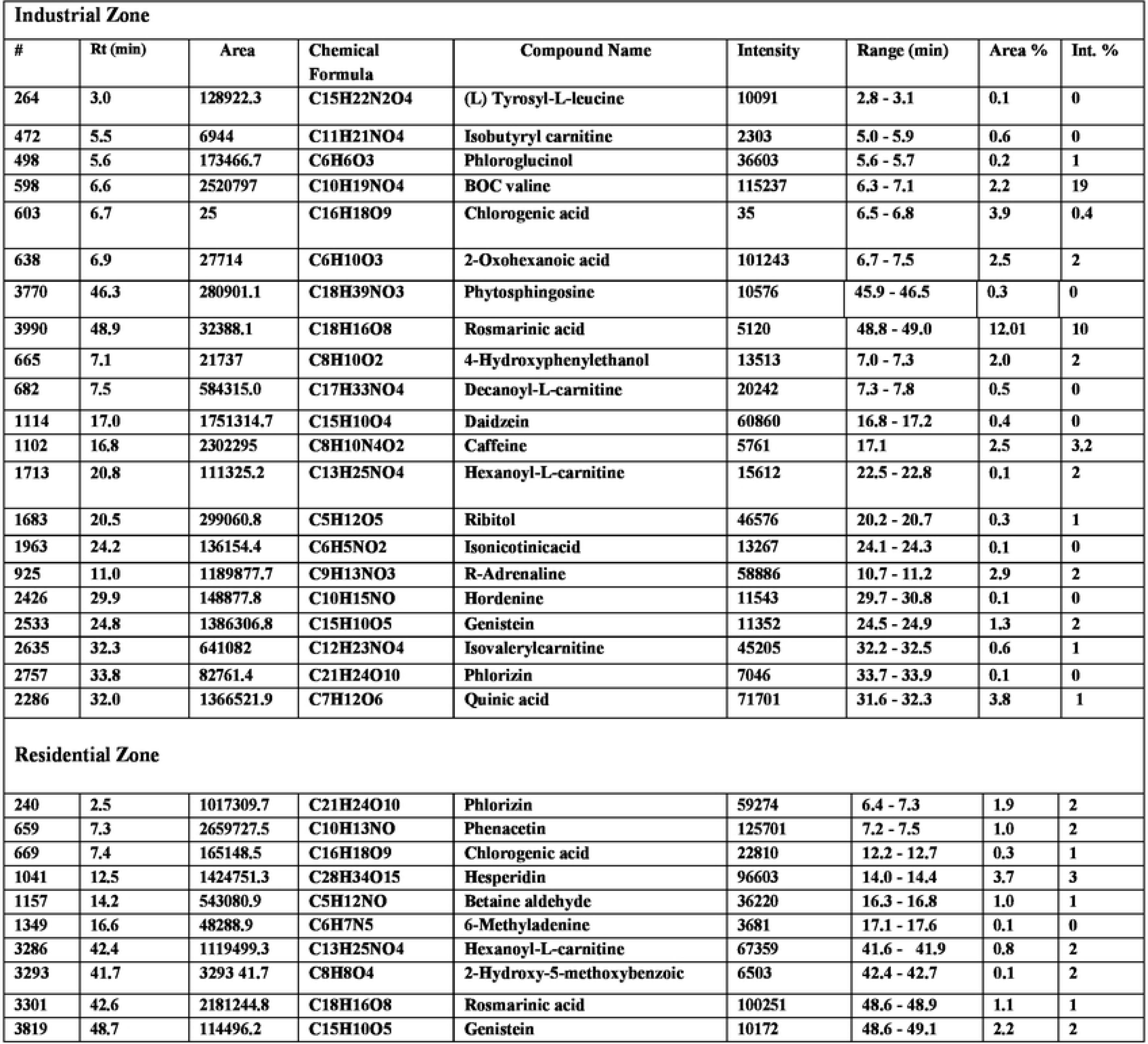
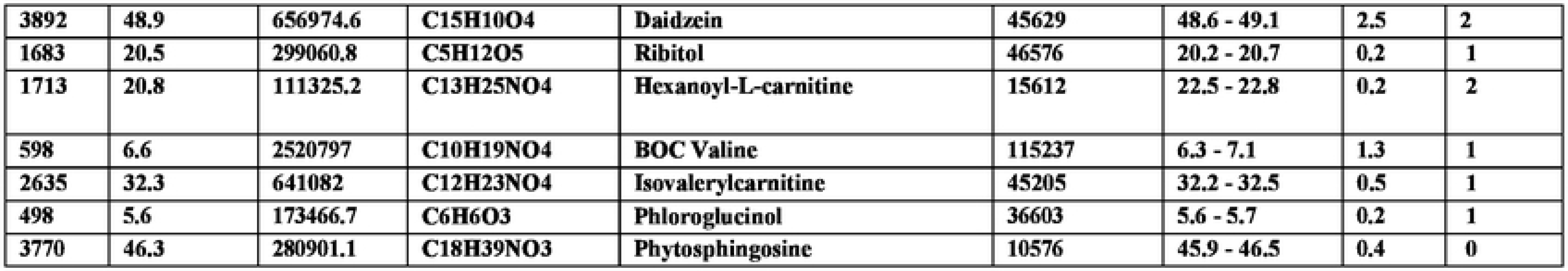
HPLC-MS analysis of methanolic extracts of *Bougainvillea glabra* leaves growing in the industrial and residential zones.

| Industrial Zone |  |  |  |  |  |  |  |  |
| --- | --- | --- | --- | --- | --- | --- | --- | --- |
| # | Rt (min) | Area | Chemical Formula | Compound Name | Intensity | Range (min) | Area % | Int. % |
| 264 | 3.0 | 128922.3 | C15H22N2O4 | (L) Tyrosyl-L-leucine | 10091 | 2.8 - 3.1 | 0.1 | 0 |
| 472 | 5.5 | 6944 | C11H21NO4 | Isobutyryl carnitine | 2303 | 5.0 - 5.9 | 0.6 | 0 |
| 498 | 5.6 | 173466.7 | C6H6O3 | Phloroglucinol | 36603 | 5.6 - 5.7 | 0.2 | 1 |
| 598 | 6.6 | 2520797 | C10H19NO4 | BOC valine | 115237 | 6.3 - 7.1 | 2.2 | 19 |
| 603 | 6.7 | 25 | C16H18O9 | Chlorogenic acid | 35 | 6.5 - 6.8 | 3.9 | 0.4 |
| 638 | 6.9 | 27714 | C6H10O3 | 2-Oxohexanoic acid | 101243 | 6.7 - 7.5 | 2.5 | 2 |
| 3770 | 46.3 | 280901.1 | C18H39NO3 | Phytosphingosine | 10576 | 45.9 - 46.5 | 0.3 | 0 |
| 3990 | 48.9 | 32388.1 | C18H16O8 | Rosmarinic acid | 5120 | 48.8 - 49.0 | 12.01 | 10 |
| 665 | 7.1 | 21737 | C8H10O2 | 4-Hydroxyphenylethanol | 13513 | 7.0 - 7.3 | 2.0 | 2 |
| 682 | 7.5 | 584315.0 | C17H33NO4 | Decanoyl-L-carnitine | 20242 | 7.3 - 7.8 | 0.5 | 0 |
| 1114 | 17.0 | 1751314.7 | C15H10O4 | Daidzein | 60860 | 16.8 - 17.2 | 0.4 | 0 |
| 1102 | 16.8 | 2302295 | C8H10N4O2 | Caffeine | 5761 | 17.1 | 2.5 | 3.2 |
| 1713 | 20.8 | 111325.2 | C13H25NO4 | Hexanoyl-L-carnitine | 15612 | 22.5 - 22.8 | 0.1 | 2 |
| 1683 | 20.5 | 299060.8 | C5H12O5 | Ribitol | 46576 | 20.2 - 20.7 | 0.3 | 1 |
| 1963 | 24.2 | 136154.4 | C6H5NO2 | Isonicotinicacid | 13267 | 24.1 - 24.3 | 0.1 | 0 |
| 925 | 11.0 | 1189877.7 | C9H13NO3 | R-Adrenaline | 58886 | 10.7 - 11.2 | 2.9 | 2 |
| 2426 | 29.9 | 148877.8 | C10H15NO | Hordenine | 11543 | 29.7 - 30.8 | 0.1 | 0 |
| 2533 | 24.8 | 1386306.8 | C15H10O5 | Genistein | 11352 | 24.5 - 24.9 | 1.3 | 2 |
| 2635 | 32.3 | 641082 | C12H23NO4 | Isovalerylcarnitine | 45205 | 32.2 - 32.5 | 0.6 | 1 |
| 2757 | 33.8 | 82761.4 | C21H24O10 | Phlorizin | 7046 | 33.7 - 33.9 | 0.1 | 0 |
| 2286 | 32.0 | 1366521.9 | C7H12O6 | Quinic acid | 71701 | 31.6 - 32.3 | 3.8 | 1 |
| Residential Zone |  |  |  |  |  |  |  |  |
| 240 | 2.5 | 1017309.7 | C21H24O10 | Phlorizin | 59274 | 6.4 - 7.3 | 1.9 | 2 |
| 659 | 7.3 | 2659727.5 | C10H13NO | Phenacetin | 125701 | 7.2 - 7.5 | 1.0 | 2 |
| 669 | 7.4 | 165148.5 | C16H18O9 | Chlorogenic acid | 22810 | 12.2 - 12.7 | 0.3 | 1 |
| 1041 | 12.5 | 1424751.3 | C28H34O15 | Hesperidin | 96603 | 14.0 - 14.4 | 3.7 | 3 |
| 1157 | 14.2 | 543080.9 | C5H12NO | Betaine aldehyde | 36220 | 16.3 - 16.8 | 1.0 | 1 |
| 1349 | 16.6 | 48288.9 | C6H7N5 | 6-Methyladenine | 3681 | 17.1 - 17.6 | 0.1 | 0 |
| 3286 | 42.4 | 1119499.3 | C13H25NO4 | Hexanoyl-L-carnitine | 67359 | 41.6 - 41.9 | 0.8 | 2 |
| 3293 | 41.7 | 3293 41.7 | C8H8O4 | 2-Hydroxy-5-methoxybenzoic | 6503 | 42.4 - 42.7 | 0.1 | 2 |
| 3301 | 42.6 | 2181244.8 | C18H16O8 | Rosmarinic acid | 100251 | 48.6 - 48.9 | 1.1 | 1 |
| 3819 | 48.7 | 114496.2 | C15H10O5 | Genistein | 10172 | 48.6 - 49.1 | 2.2 | 2 |
| 3892 | 48.9 | 656974.6 | C15H10O4 | Daidzein | 45629 | 48.6 - 49.1 | 2.5 | 2 |
| 1683 | 20.5 | 299060.8 | C5H12O5 | Ribitol | 46576 | 20.2 - 20.7 | 0.2 | 1 |
| 1713 | 20.8 | 111325.2 | C13H25NO4 | Hexanoyl-L-carnitine | 15612 | 22.5 - 22.8 | 0.2 | 2 |
| 598 | 6.6 | 2520797 | C10H19NO4 | BOC Valine | 115237 | 6.3 - 7.1 | 1.3 | 1 |
| 2635 | 32.3 | 641082 | C12H23NO4 | Isovalerylcarnitine | 45205 | 32.2 - 32.5 | 0.5 | 1 |
| 498 | 5.6 | 173466.7 | C6H6O3 | Phloroglucinol | 36603 | 5.6 - 5.7 | 0.2 | 1 |
| 3770 | 46.3 | 280901.1 | C18H39NO3 | Phytosphingosine | 10576 | 45.9 - 46.5 | 0.4 | 0 |

**Fig. 3.**
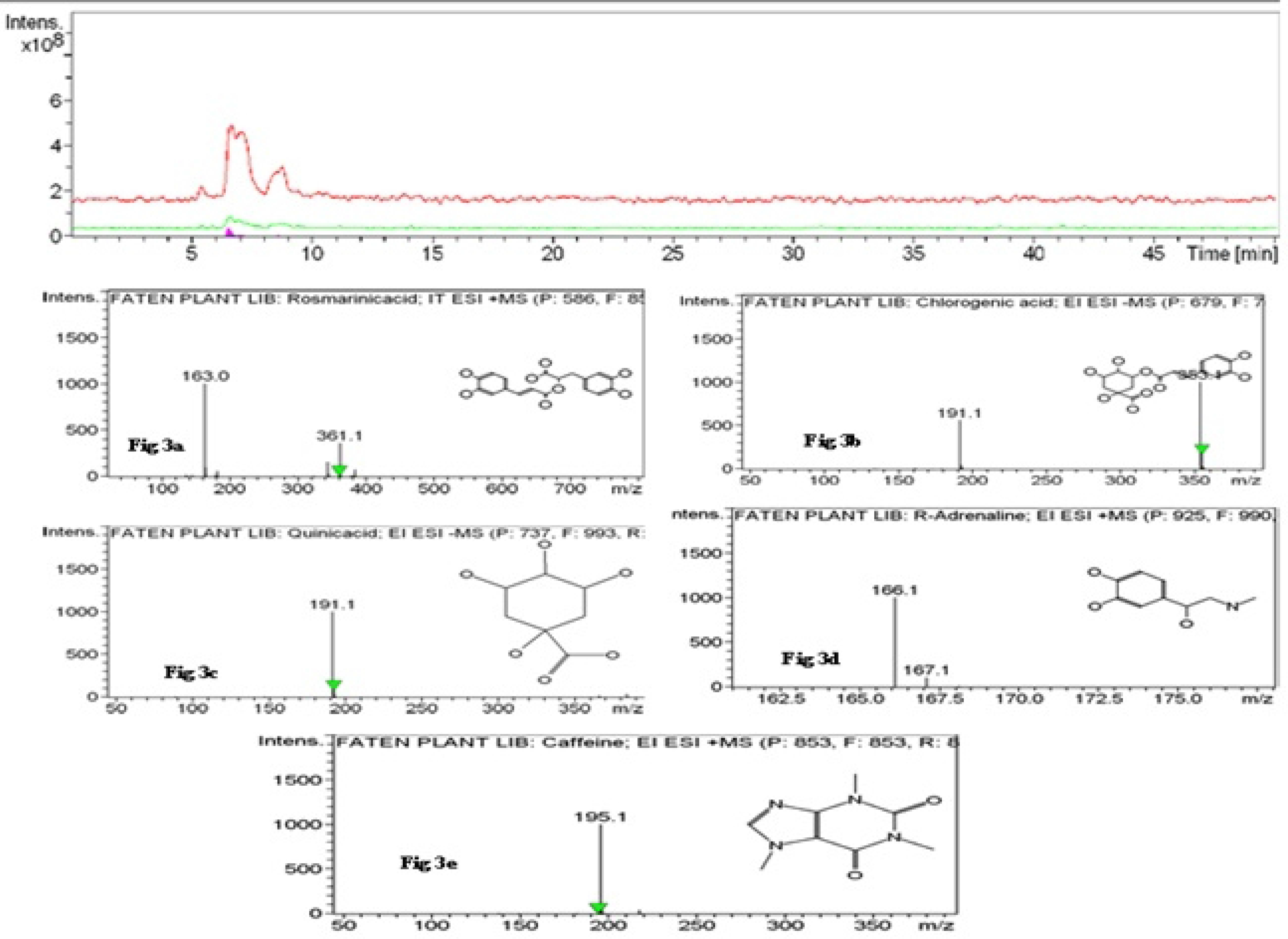
HPLC-MS analysis of methanolic extracts of *Bougainvillea glabra* leaves growing in the industrial zone.

The HPLC-MS analysis of *Bougainvillea glabra* from the residential area (**Table 3**, **Fig. 4**) revealed seventeen compounds all identified as flavonoids, four of which were major peaks with retention times of 5 min and over: hesperidin (Rt 12.5 min), peak area percentage 3.7%; daidzein (Rt 48.9 min), peak area 2.5%; genistein (Rt 48.7 min), peak area 2.2%, and phlorizin (Rt 2.5 min), peak area 1.9%.

**Fig. 4.**
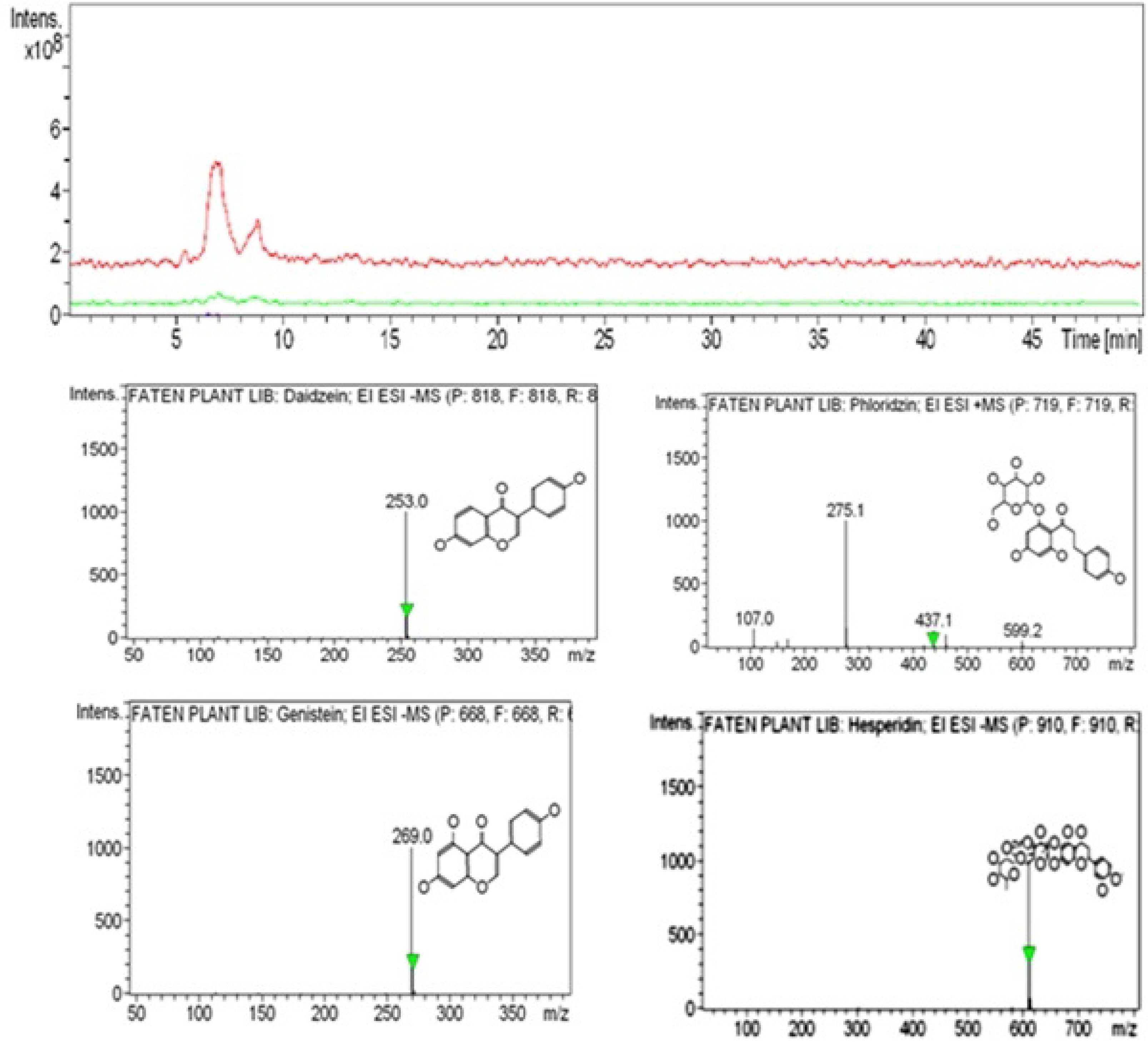
HPLC-MS analysis of methanolic extracts of *Bogainvillea glabra* leaves growing in the residential zone

Taken together, the total phenolics and flavonoids of *B. glabra* were significantly higher in the industrial zone than the residential zone (P<0.05; **Table 4**).

**Table 4.** Determination of total phenolic and flavonoid contents in *Bougainvillea glabra* leaf extracts. Values are means ± SD from three replicates.

|  | Industrial Zone | Residential Zone | Sign. |
| --- | --- | --- | --- |
| Total phenolics | 61.72 ± 0.70 GAE mg/100g | 45.82 ± 0.50 GAE mg/100g | P<0.05 |
| Total flavonoids | 233.53 ± 16.10 QE mg/100g | 212.23 ± 9.05 QE mg/100g | P<0.05 |

### 3.4 Total suspended particulates (TSP, PM_10_) and gases in the industrial and residential zones

**Table 5** shows that there were significantly higher concentrations of TSPs, PM_10_, and gases at the industrial site compared to the residential zone.

**Table 5.** Means of TSP-PM_10_ and gases in the industrial and residential zones during March and April 2017/2018 (mg/m^3^). Values are means ± SD from three replicates.

| Item | Study Zone and Value |  | Sign. |
| --- | --- | --- | --- |
| TSP | Industrial | 7.06 ± 2.3 | P<0.05 |
|  | Residential | 0.40 ± 0 |  |
| PM<sub>10</sub> | Industrial | 15.72 ± 6.5 | P<0.05 |
|  | Residential | 0.0 ± 0.0 |  |
| NO | Industrial | 9.66 ± 5.0 | P<0.05 |
|  | Residential | 0.0 ± 0.0 |  |
| NO<sub>2</sub> | Industrial | 0.12 ± 0.04 | P<0.05 |
|  | Residential | 0.04 ± 0.0 |  |
| NO<sub>x</sub> | Industrial | 68.43 ± 1.66 | P<0.05 |
|  | Residential | 0.0 ± 0.0 |  |
| SO<sub>2</sub> | Industrial | 16.76 ± 12.74 | P<0.05 |
|  | Residential | 0.05 ± 0.0 |  |
| CO | Industrial | 42.16 ± 24.1 | P<0.05 |
|  | Residential | 0.0 ± 0.0 |  |
| CO<sub>2</sub> | Industrial | 473.00 ± 35.5 | P<0.05 |
|  | Residential | 0.01 ± 0.0 |  |

### 3.5 Groundwater analysis

Table 6 shows that turbidity, total distilled solids (TDS), pH, chlorides, fluorides, sulfates, hardness, calcium, magnesium, silicon dioxide, zinc, and selenium were all significantly higher in water from the industrial zone than the residential zone (all P<0.05) and potassium and barium were significantly lower in water from the industrial zone than the residential zone (all P<0.05).

**Table 6.**
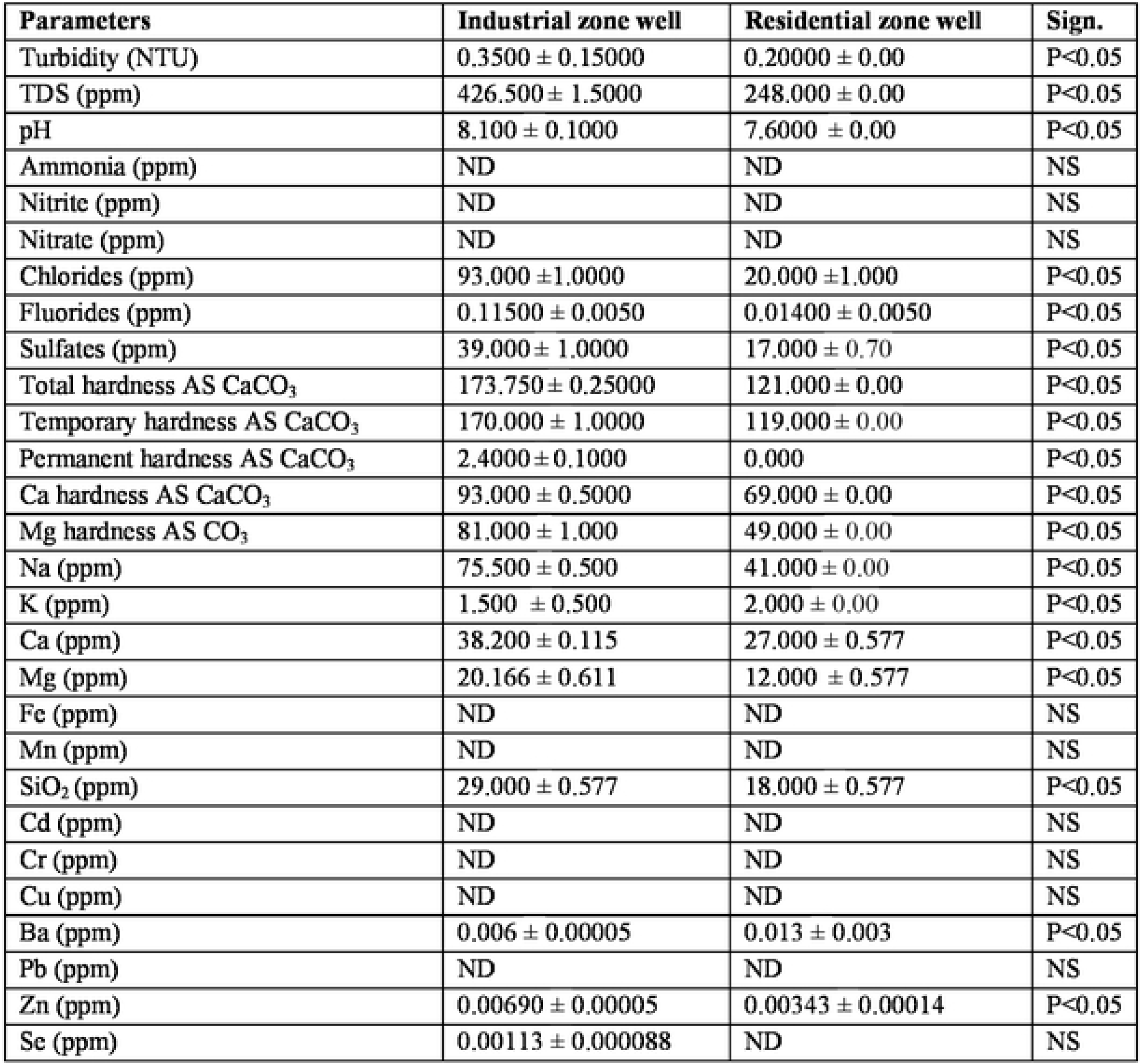
Water analysis from wells in the residential and industrial zones. ND=not detected, NTU=nephelometric turbidity unit, NS=not significant, TDS=total distilled solids. Values are means ± SD from three replicates.

| Parameters | Industrial zone well | Residential zone well | Sign. |
| --- | --- | --- | --- |
| Turbidity (NTU) | 0.3500 ± 0.15000 | 0.20000 ± 0.00 | P<0.05 |
| TDS (ppm) | 426.500 ± 1.5000 | 248.000 ± 0.00 | P<0.05 |
| pH | 8.100 ± 0.1000 | 7.6000 ± 0.00 | P<0.05 |
| Ammonia (ppm) | ND | ND | NS |
| Nitrite (ppm) | ND | ND | NS |
| Nitrate (ppm) | ND | ND | NS |
| Chlorides (ppm) | 93.000 ± 1.0000 | 20.000 ± 1.000 | P<0.05 |
| Fluorides (ppm) | 0.11500 ± 0.0050 | 0.01400 ± 0.0050 | P<0.05 |
| Sulfates (ppm) | 39.000 ± 1.0000 | 17.000 ± 0.70 | P<0.05 |
| Total hardness AS CaCO<sub>3</sub> | 173.750 ± 0.25000 | 121.000 ± 0.00 | P<0.05 |
| Temporary hardness AS CaCO<sub>3</sub> | 170.000 ± 1.0000 | 119.000 ± 0.00 | P<0.05 |
| Permanent hardness AS CaCO<sub>3</sub> | 2.4000 ± 0.1000 | 0.000 | P<0.05 |
| Ca hardness AS CaCO<sub>3</sub> | 93.000 ± 0.5000 | 69.000 ± 0.00 | P<0.05 |
| Mg hardness AS CO<sub>3</sub> | 81.000 ± 1.000 | 49.000 ± 0.00 | P<0.05 |
| Na (ppm) | 75.500 ± 0.500 | 41.000 ± 0.00 | P<0.05 |
| K (ppm) | 1.500 ± 0.500 | 2.000 ± 0.00 | P<0.05 |
| Ca (ppm) | 38.200 ± 0.115 | 27.000 ± 0.577 | P<0.05 |
| Mg (ppm) | 20.166 ± 0.611 | 12.000 ± 0.577 | P<0.05 |
| Fe (ppm) | ND | ND | NS |
| Mn (ppm) | ND | ND | NS |
| SiO<sub>2</sub> (ppm) | 29.000 ± 0.577 | 18.000 ± 0.577 | P<0.05 |
| Cd (ppm) | ND | ND | NS |
| Cr (ppm) | ND | ND | NS |
| Cu (ppm) | ND | ND | NS |
| Ba (ppm) | 0.006 ± 0.00005 | 0.013 ± 0.003 | P<0.05 |
| Pb (ppm) | ND | ND | NS |
| Zn (ppm) | 0.00690 ± 0.00005 | 0.00343 ± 0.00014 | P<0.05 |
| Se (ppm) | 0.00113 ± 0.000088 | ND | NS |

### 3.6 Chemical analysis of soil samples

Heavy metal concentrations in the soil samples from industrial and residential zones are listed in Table 7. All metals apart from cadmium were present at significantly higher levels in the samples obtained from the industrial area than the residential area (all P<0.05).

**Table 7.** Means of heavy metals concentrations in soil samples (ppm) collected from industrial and residential zones. Values are means ± SD from three replicates.

| Metal | Industrial Zone | Residential<br>Zone | Sign. |
| --- | --- | --- | --- |
| Fe | 8750.00 ± 21.6 | 5559.0 ± 0.57 | P<0.05 |
| Cu | 2.08 ± 0.20 | 1.37 ± 0.008 | P<0.05 |
| Cd | 0 | 0 | NS |
| Zn | 12.50 ± 0.0 | 0.28 ± 0.005 | P<0.05 |
| Mn | 33.33 ± 4.16 | 19.71 ± 0.005 | P<0.05 |
| Pb | 4.57 ± 0.15 | 0.48 ± 0.005 | P<0.05 |
| Ni | 20.48 ± 2.02 | 0.500 ± 0.005 | P<0.05 |
| Cr | 30.29 ± 2.2 | 23.50 ± 0.005 | P<0.05 |
| Co | 0.39 ± 0.013 | 1.25 ± 0.005 | P<0.05 |

## 4. DISCUSSION

Sadat City, located north of Cairo, is characterized by extreme aridity, a long hot summer, and short warm winter, which greatly influences the hydrological properties of the drainage basins in the area [25, 26]. Rapid industrialization is causing health hazards, and the effect of industrialization on human health is a major public health concern [27].

Morphological examination of plants growing in industrial and residential zones revealed marked differences in the leaves in different areas. [28] Reported that environmental stresses negatively influence growth, productivity, and trigger a series of morphological, physiological, biochemical and molecular changes in plants. Dust deposition on leaf surfaces may also reduce chlorophyll synthesis due to a shading effect [29]. The resulting changes, either morphological or physiological, can be used as bioindicators of the environmental state [30].

Flavonoids function as stress indicators because they accumulate at high levels in many plant tissues in response to a wide range of biotic and abiotic signals [2]. Flavonoids comprise the large and common group of plant phenolics with more than 5000 different described flavonoids in six major subclasses [31]. Plants may alter their secondary metabolite synthesis, production, secretion and storage when subjected to the abiotic stress factors [32]. The results obtained in the present study revealed that pollution in the industrial area was associated with elevations in total phenolics and flavonoids compared with samples of plants from non-polluted, residential sites. The elevations in total phenolics and flavonoids may have acted as a stress defense mechanism in plants against these environmental pollutants.

HPLC-MS profiles of the studied plant samples from the industrial zone revealed the presence of 21 compounds, five of which were present as major peaks belonging to flavonoid and phenolic compounds. R-adrenaline was detected in plants growing in the industrial zone, which may due to environmental stress; indeed, [33] reported that adrenaline is known to protect against oxidation by flavonoids. [32] Reported that flavonoids comprise a large and common group of plant phenolics, and flavonoids commonly increases in response to unfavorable conditions. By contrast, residential zone plant extracts showed the presence of 17 compounds, four of which were major flavonoid and phenolic peaks. Therefore, total phenolics and flavonoids highly represented in plants in the industrial zone may be a result of stress-inducing pollution, and reported previously [34]. Phenolic compounds (flavonoids) function as stress indicators because they accumulate at high levels in many plant tissues in response to a wide range of biotic and abiotic signals [35 –37], enhancing the scavenging of free radicals [38]. Our results are also consistent with [39], who observed higher quercetin (phenolic flavonoid) content in samples from a polluted site, suggesting that the increase could be associated with the defensive role of flavonoids under conditions of environmental stress. Flavonoids are a good indicator of environmental contamination, especially of O_3_ pollution [40].

The total concentrations of TSPs in water in the industrial zone were higher than the US Environmental Protection Agency’s prescribed standards for air quality, presumably as a result of emissions from local industrial activity such as steel factories in this area. Generally, the outlet values of TSPs in samples were less than legally permitted values (EEAA (Egyptian Ministry for the Environment) Law 4/1994; 9/2009), 10 mg/m^3^ in the work environment. Similarly, the PM_10_ were higher in the industrial zone and only present at trace levels in the residential zone, but the outlet PM_10_ values in all samples were generally lower than required by law (4/1994; 9/2009). The higher SO_2_ values at the industrial site may have environmental consequences; NOx dissolves in cells and gives rise to nitrite ions (NO_2_ which is toxic at high concentrations) and nitrate ions (NO_3_) that enter into nitrogen metabolism as if they had been absorbed through the roots. In some cases, exposure to pollutant gases, particularly SO_2_, causes stomata closure, which protects the leaf against the further entry of the pollutant but also curtails photosynthesis [1].

The high level of contamination with iron and other heavy metals in plants and soil in the industrial zone could have originated from metal-containing waste from the steel industry. In general, the heavy metals concentrations were lower than allowed by law (4/1994; 9/2009). Cadmium and copper were not detected in either zone, perhaps because they were not used as raw materials in any of the industries. The U.S. Environmental Protection Agency [41] regulates nine trace elements for land-applied sewage sludge (As, Cd, Cu, Pb, Hg, Mo, Ni, Se, and Zn), with six of these elements (Cu, Ni, Zn, Cd, Pb, and Se) considered to be phytotoxic (The quality of all groundwater samples was compatible with the Egyptian standards for drinking water and hosing.

Lead, a toxic heavy metal [42] was recorded in plants in the industrial zone but not the residential zone. Possible sources of lead are batteries, chemical substances from photograph processing, lead-based paints, and lead pipes deposited in the landfill. The high levels of Zn in the plants, water, and soil in the industrial zone compared to the residential zone could be attributable to metal-containing waste in the industrial zone, which might have leached into the underlying soil to be absorbed by plants [43].

## 5. CONCLUSIONS

This study compares the presence and levels of pollutants between residential and industrial areas of a city and their possible effects on plants. Pollution is becoming an ever-growing global environmental problem. Industrialization, population growth, and the associated increase in energy demands have resulted in a profound deterioration of the environmental quality in developing countries like Egypt. Pollution comes from many different sources such as factories, automobiles, and even wind-blown dust. Pollution is not harmful only to plants but to all living organisms. Plants are very good bioindicators of pollution and may be used as pollution biomarkers.

## AUTHOR CONTRIBUTIONS

MFA designed the study, conducted the experiments, and wrote the manuscript.

